# eOceans dive-logs for science and conservation: a case study of sharks in Thailand

**DOI:** 10.1101/296160

**Authors:** C. A. Ward-Paige, A. Westell, B. Sing

## Abstract

Many marine animals around the world are threatened by a variety of anthropogenic activities, yet there is often a paucity of data to monitor patterns in abundance and distribution or to evaluate human interventions. The new citizen science program eOceans helps to fill this gap by gathering observations of various marine animals from worldwide ocean explorers. In 2012, a dedicated Thailand-wide census of sharks, and other animals, began as collaboration between eOceans scientists and the dive tourism industry. Using the observations from 9,524 dives (9,357 hours underwater) logged by >169 divers on 153 sites, we describe the spatial and temporal patterns of sharks in coastal Thailand. A total of 12 shark species were encountered, most commonly (67%) as individuals, and were observed on 11% of all dives, on 59% of sites, in all months and years. The two most frequently encountered species were blacktip reef (*Carcharhinus melanopterus*) and leopard sharks (*Stegostoma fasciatum*). Many species had peak encounter rates in summer, but aggregated in various seasons in different years. Mating events and nursery sites were observed rarely, and only for blacktip reef and whitetip reef (*Triaenodon obesus*) sharks. These results could be of value to species-or region-specific biologists, ecologists and fisheries scientists, as well as to managers and policy makers that could use the findings to monitor future trends and prioritize conservation strategies. Moreover, this study highlights the value that collaborative eOceans citizen science projects could have in support of marine science, management and conservation efforts worldwide.

## I. INTRODUCTION

Many marine animal populations have been, or continue to be threatened by anthropogenic impacts (Lotze et al., 2011), and sharks are among the most threatened animal groups, affected by fishing, habitat loss, and climate change (Dulvy et al., 2014; Oliver et al., 2015; Worm et al., 2013). Coastal sharks, including reef sharks, have been repeatedly shown to have declined long ago, pre-dating fishing records (Nance et al., 2011) and modern ecological assessments (Ferretti et al., 2010, 2008; Sandin et al., 2008; Ward-Paige et al., 2010b). This lack of data and protracted history of overexploitation means that historic population trends are often missing and, since fishing and other anthropogenic activities continue to affect populations, establishing rates of population change are challenging to attain and many species may be more vulnerable than previously thought (Osgood and Baum, 2015). As well, most shark research tends to occur in areas where sharks are still relatively abundant, leaving many heavily populated and overexploited coastlines undocumented by scientific observations, and therefore even contemporary baselines are not being established for monitoring into the future.

Typically, scientific population censuses of sharks are made by utilizing fisheries-dependent data, such as catch or bycatch data (e.g., Baum et al., 2003; Carlson et al., 2012; Ferretti et al., 2008) or fisheries independent data that may still involve high mortality sampling techniques, such as with gillnets (Froeschke et al., 2010; e.g., Ward-Paige et al., 2015) or trawls (Ferretti et al., 2010). Some aim to lower mortality during sampling (Hammerschlag and Sulikowski, 2011), especially where tags for mark-recapture or tracking studies are being used (Speed et al., 2011). However, in many areas where lethal sampling is illegal or unacceptable, such as on coral reefs, coastal areas near tourist sites, and marine protected areas, non-lethal sampling is often sought. In these cases, baited remote underwater visual censuses (BRUV; Brooks et al., 2013; Colton and Swearer, 2010) and underwater visual censuses (UVC) done by scientific scuba divers (Robbins et al., 2006; Ward-Paige et al., 2010a) or trained volunteer divers (e.g., transects, Reef Life Survey (reeflifesurvey.org)) have been used. Similar to lethal sampling techniques, however, those focusing on sharks also typically take place where sharks are relatively abundant. In other areas, where sharks are rare and BRUVs or UVC are used to survey other animal groups (e.g., corals, reef fish), some sharks may be encountered, but often occur too infrequently to describe populations.

Around the world, individuals with a range of experiences, are undertaking marine tourist activities, making millions of observations on a daily basis, and reporting what they observe to personal log books, or to various science or non-profit organizations. Since the mid-2000s, many citizen science census platforms have been launched to collect these observations. Some have provided context for understanding the value and limitations of these data (Vianna et al., 2014; Ward-Paige et al., 2010a; Ward-Paige and Lotze, 2011), while others have provided important insights on shark ecology, including reproductive seasonality, fisheries interactions, and movements (Bansemer and Bennett, 2010, 2008; Whitney et al., 2011), and population status (Theberge and Dearden, 2006; Vianna et al., 2014; Ward-Paige et al., 2013, 2010b; White et al., 2015). These projects range from being species specific (e.g., whitetip reef, *Triaenodon obesus*, Whitney et al., 2011), which collect presence only data, to exhaustive checklists with abundance (e.g., all fish species; Reef Environmental Education Foundation (REEF.org)). Each has advantages and disadvantages. For example, species-specific projects lack non-target species and observations where no target species were observed (i.e., zeros), but can be rolled out in a target area relatively quickly and garner focused attention on the target species. Checklists that include all species, on the other hand, require extensive training and time to roll out, but get zeros, which is valuable for monitoring trends through time and space.

Because sharks are mobile and can move around and between sites there is need for high effort for detection. This is especially true for sharks that are rare, depleted, or seasonal, and may only infrequently visit sites. Correct identification to species level, however, can be a challenge for reasons ranging from encounters that are too brief to a lack of training (e.g., Brunnschweiler, 2009). However, compared to other animal groups that are regularly studied by citizen observers (e.g., eBird, Sullivan et al., 2009), the characteristics of sharks make them relatively easy to identify. Birds, for example, have checklists containing 9,000-10,000 species, while there are only about 500 shark species in total (Dulvy et al., 2014), and fewer live within the depth range of divers. As well, some, especially reef-associated sharks (Osgood and Baum, 2015), have some site fidelity and are repeatedly observed and photographed by divers in-situ, which can further increase identification accuracy. Additionally, recreational divers that roam a site are better able to collect data on low density and rare fishes than many scientific divers, who use predefined transects, and they cover a bigger area and diverse habitat types (Ward-Paige and Lotze, 2011). Therefore, with appropriate caution, using recreational divers’ observations can be ideal to increase observation effort data, without the requirement of extensive training or photographs.

eOceans (www.eOceans.co) is an umbrella program that hosts various marine-focused citizen science projects (e.g., previously eShark, eManta, Global Marine Conservation Assessment). It provides an online platform where all marine explorers are invited to enter either (i) ‘snapshot summaries’ of past observations for hypothesis driven research questions, such as to describe the distribution and human use patterns of manta rays (Ward-Paige et al., 2013); or (ii) ‘event-based reports’ of each ocean experience (e.g., every dive) for ongoing, high-resolution monitoring of animal populations at specific sites (current study). The platform was developed using insights gained from years of in-depth investigations (by author CAWP) of the value and limitations of recreational divers’ observations for providing shark and ray observations (Ward-Paige et al., 2014, 2010a, 2010b; Ward-Paige and Lotze, 2011). Although the primary focus was initially to collect data on sharks (and was previously called eShark), it also collects observations of rays, turtles, seahorses, jellyfish, whales, dolphins, seals, and marine debris, depending on location. These additions were found to increase participation, reporting of zeros, and the versatility of the dataset. See further details behind the development, implementation and communication strategies of eOceans in Hind-Ozan et al. (2017).

In 2012, the Thailand tourism industry, lead by the local non-profit organization ‘Shark Guardian’ (sharkguardian.org), launched a nation-wide concentrated dive census for eOceans. Through this, invitations were sent out extensively to SCUBA diving companies, clubs, and divers across the country, particularly in coastal tourist regions, to participate by submitting observations from every dive. Here, we use these reports to describe spatial and temporal patterns of surveyor effort and shark populations in Thailand. These findings demonstrate the immense potential of eOceans as a community driven (i.e., bottom-up, see Roelfsema et al., 2016) marine citizen science project, for providing relatively high-resolution temporal information at the site and regional scale. They may also assist species-specific or region-specific biologists and ecologists, as well as managers and policy makers to prioritize scientific investigations and conservation strategies of sharks in Thailand. As well, additional eOceans nation-wide projects (currently in Fiji, Indonesia, and South Africa) could immensely expand our knowledge of some coastal shark populations, and other species, which could help focus scientific investigations and conservation tactics on a regional and global scale.

## II METHODS

The eOceans platform has an online form that collects event-based (i.e., every dive) information from various ocean explorers (divers, snorkelers, fishers, etc.). Data collected from scuba divers (the focus of this study) included contact email, dive experience (number of dives in life), dive location (country, area, GPS coordinates and/or site name,), dive date, the use of attractant (e.g., chumming, baiting) or spearfishing, and the presence or absence of jellyfish, seahorses, turtles, mammals, and litter, as well as the number of sharks and rays by species. For each, ‘unknown’ is offered as an option. Observations of sharks actively involved in mating or possible nursery locations (i.e., where many small individuals were observed together), were also solicited. Since 2007, there have been two versions of the form. Version 2 (V2) replaced version 1 (V1) in November 2015. Both had similar objectives and questions, but differed slightly. V1 asked for site depth and habitat type, which was redundant for dives occurring at the same site, and were therefore included as a descriptor within the site dropdown list in V2. V1 had photos of different sharks to select from that were grouped by like species (e.g., nurse sharks with a dropdown menu for tawny nurse shark), whereas V2 had a dropdown list of the most commonly sighted species, with room to enter other species, and no photos were included.

In 2012, members of the recreational dive industry in Thailand, led by the non-profit group ‘Shark Guardian’ (www.sharkguardian.org), committed to submitting their daily dive observations. Dive shops typically visit 10-20 different dive sites, up to three per day, which are scheduled by the day of the week. Site schedules usually only changed according to the weather, not the presence or absence of certain species (e.g., they do not change sites to target or avoid sharks). Divers in Thailand had the option of reporting observations directly online, or by recording dives and observations into community logbooks (e.g., at a dive shop) immediately following a dive. This commenced the first-ever, nation-wide census of sharks in Thailand. Dive shops, dive guides, their clients, and other recreational divers, participated by reporting daily observations. Shark Guardian recruited members by delivering a brief (20 minute) presentation to interested participants, which included reasons for the study (e.g., vulnerability of sharks), how to correctly identify shark species, and where to report observations (See presentation outline here: http://www.sharkguardian.org/the-shark-guardian-presentation/).

For the current study, only events that took place from 2012-2016 by scuba diving and snorkeling activities were included (omitting surfing, fishing, and other activities because they were too uncommon). Due to limitations in the original form and species identification in general, a few reports were corrected as such: i) Tawny nurse shark (*Nebrius ferrugineus*, n = 4) was combined with “Nurse” (*Ginglymostoma cirratum*, n = 33) because they were included in the V1 online form under a single dropdown heading called ‘Nurse’, but the latter does not occur in the area, ii) Leopard shark, the local name for the zebra shark *Stegostoma fasciatum* (n = 213), was combined with Leopard shark (*Triakis semifasciata*, n = 30), which does not occur in the area but shares the same common name locally, iii) Arabian carpetshark (*Chiloscyllium arabicum*, n = 13), which does not occur in the area, was combined with the morphologically similar Grey bamboo shark (*Chiloscyllium griseum*, n = 86), which was reported on the same sites, and iv) two observations of both oceanic whitetip shark (*Carcharhinus longimanus*) and white shark (*Carcharodon carcharias*) were removed (<0.2 % of observations) as they were, after inquiry with participants, mistakes in the entry process as they were likely meant to be reported as whitetip reef sharks.

All observations were combined by species, month in each year, and site, and mapped or plotted using “lattice” plots (Sarkar, 2008) to describe spatial and temporal trends in dive effort, occurrence, sighting frequency (SF; number of times where sharks were present per the total number of dives), maximum school size, and a proxy of abundance, called ‘abundance’ from here on, which was calculated as the maximum number of sharks observed per the total number of dives.

## III. RESULTS

In Thailand, 9,524 unique dive events were reported to eOceans between 2012-2016, for a total of 9,358 hours. More than 169 individuals entered dives (note: many people provided names but not email addresses, which were considered the unique identifier and therefore an exact number of participants is not available) and average diver experience was 93 dives in life (lifetime experience), with a minimum of 1 and maximum of 5,000 dives reported. Sharks were observed on 1,053 dives (11% SF - of all dives), for a total of 2,426 sharks (including multiple sightings of the same individuals). Dive events were submitted throughout most months during the study period (Figure 1a), with increased effort in the winter and spring months (high-tourist season, October to April), reaching up to 584 dives per month. A total of 153 sites were visited with the majority (95%) of dives made in the Andaman Sea (Figure 1b), and at least one shark was observed on 90 sites (59%). Effort was distributed along both coasts, including in the Gulf of Thailand in the northeast and central-east, and in the Andaman Sea. Attractant (e.g., baiting) was used on ten dives (0.1%), on seven sites, in six different months – one of these dives had sharks present, which included 5 blacktip reef sharks. Spearfishing was used during two dives (0.02%), on two sites, and no sharks were observed during either.

**Figure 1.**
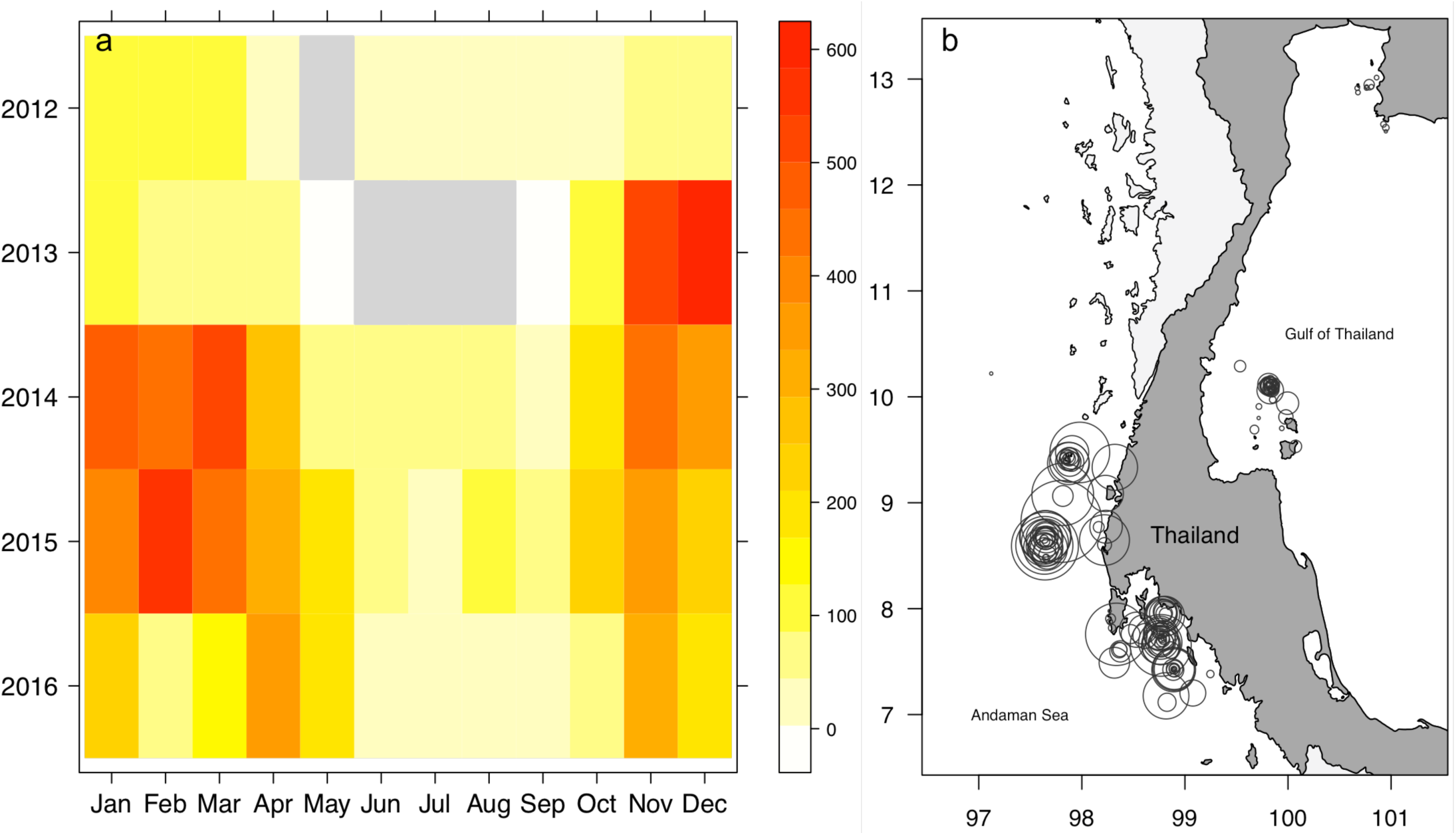
Spatial and temporal patterns of survey effort by divers in Thailand. a) Number of dives per month in each year (grey boxes indicate zero effort); and b) location of surveyed sites, where circle size is proportional to number of dives ranging from 1 to 598 dives.

Sharks actively engaged in mating or potential nursery sites were rarely reported (Figure 2). Sixteen instances of shark nurseries were reported, two of whitetip reef sharks and fourteen of blacktip reef sharks, at seven sites, in seven different months, all in 2015 and 2016. Two mating events were observed, both of blacktip reef sharks at the same site, in August 2015 and August 2016.

**Figure 2.**
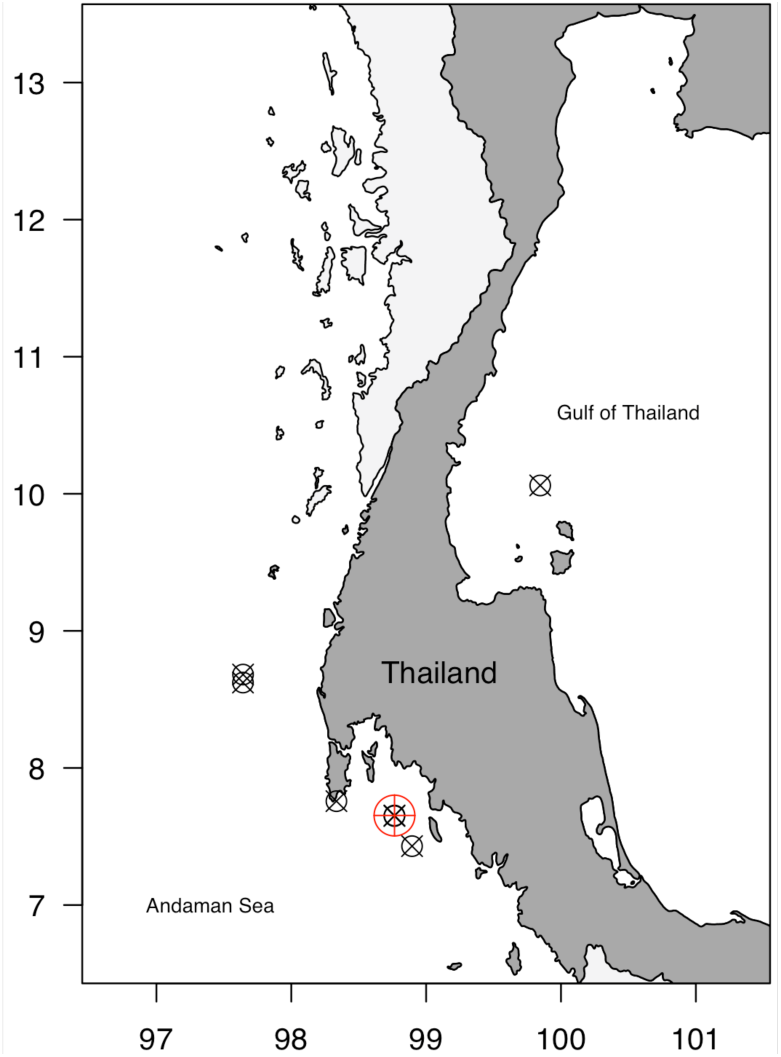
Sites with reported shark mating (red circle) and nursery (black circle) events.

A total of 12 shark species were reported, with the total number of sharks observed ranging from 8 (silvertip) to 1,282 (blacktip reef) and school sizes ranging from 1 to 45 individuals (Figure 3, Table 1). The majority (67%) of encounters were of individuals (school size = 1). Blacktip reef and leopard sharks were the most frequently encountered species, in both the most number of occurrences and the most number of individuals. Maximum school size ranged from 2 in grey reef sharks to 45 for blacktip reef sharks, and mean school size on sites where the species was observed, ranged from 1.3 in grey reef sharks to 10 for bull sharks.

**Figure 3.**
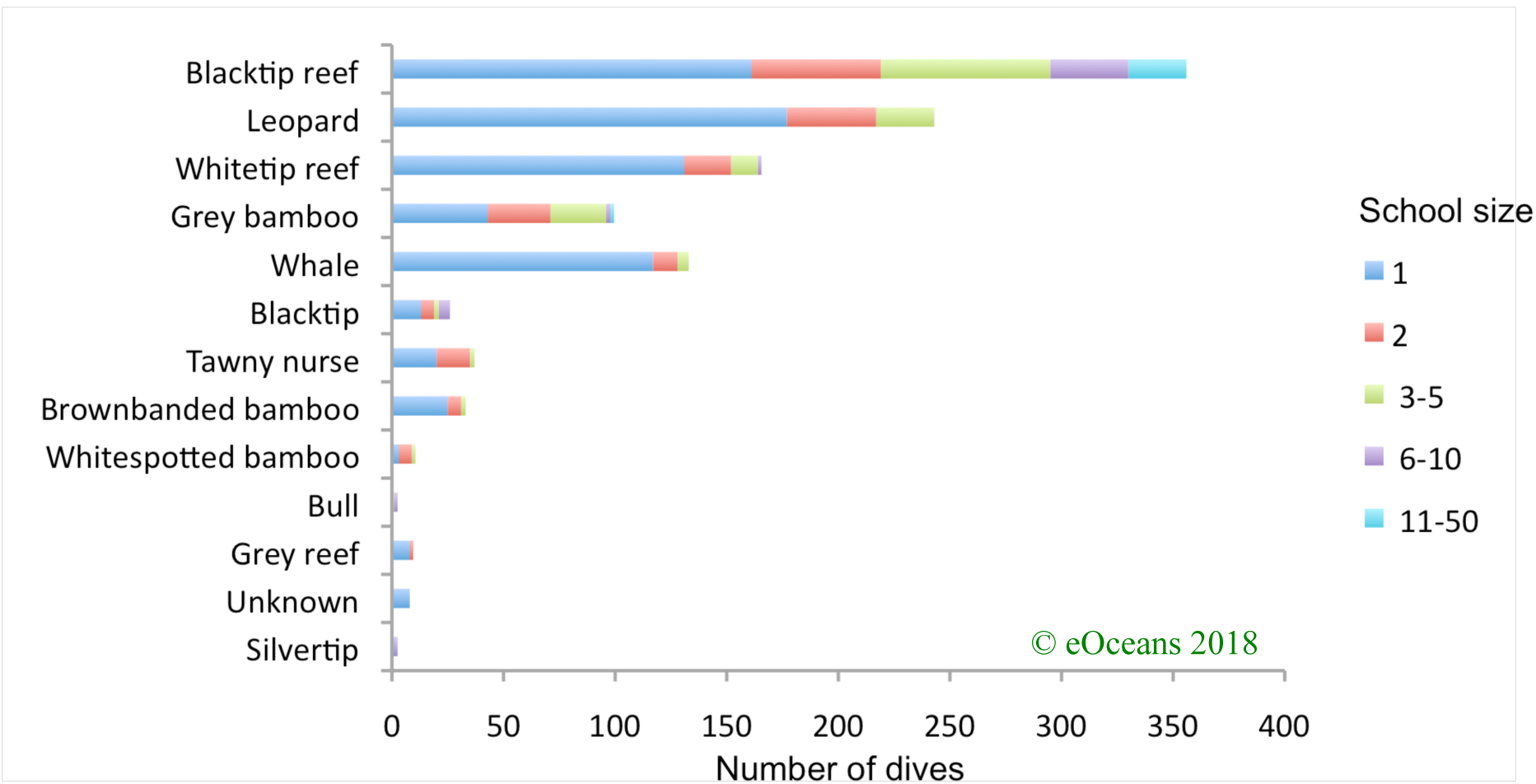
Sightings (number of dives) and school size (color coded) per species.

Sharks occurred in all months and years of the study period (coloured boxes in Figures 4 and 5; note that the scales vary between species). Patterns of occurrence, SF, and maximum school size varied by month and year for each species. Across the 56 surveyed months, monthly occurrences were highest for blacktip reef (n= 43), leopard (n = 41) and whitetip reef (n = 32) sharks (fewest white boxes in Figure 4 and 5), and all three species were observed in all years. The other nine species had more intermittent or infrequent occurrences, with brownbanded bamboo, whitespotted bamboo, bull, grey reef, and silvertip sharks only occurring in one to three years. Peaks in SF (red boxes, Figure 4) varied by season and year. Summer months (April-September) had peak SF of blacktip reef, leopard, grey bamboo, whale, tawny nurse, and whitespotted bamboo sharks, while winter months (October to March) had higher SF of blacktip and brownbanded bamboo sharks. Whitetip reef sharks had peak SF at various times of the year, and whitespotted bamboo, bull, grey reef, and silvertip sharks varied, or were too infrequently encountered to detect seasonal changes. Peak maximum school size (red boxes, Figure 5) rarely aligned with peak SF, and the months and years of peak showed very different patterns. For the three most commonly encountered species, peak maximum school size occurred at various times of the year, and changed year to year.

**Figure 4.**
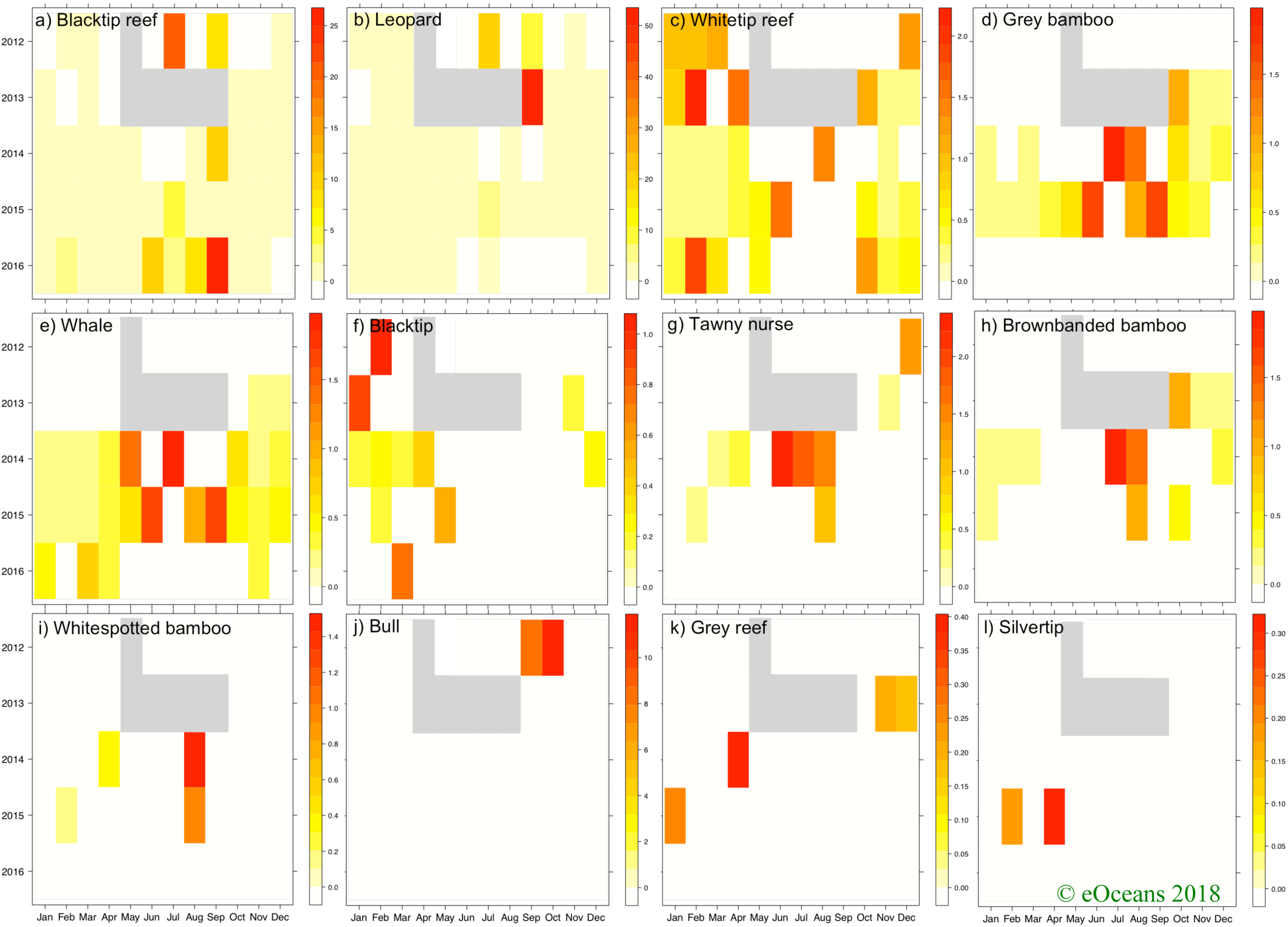
Temporal patterns of sighting frequency by species. Each scale indicates the sighting frequency - number of dives a species was observed divided by the total number of dives in that month. Grey boxes are months where sample effort was zero.

**Figure 5.**
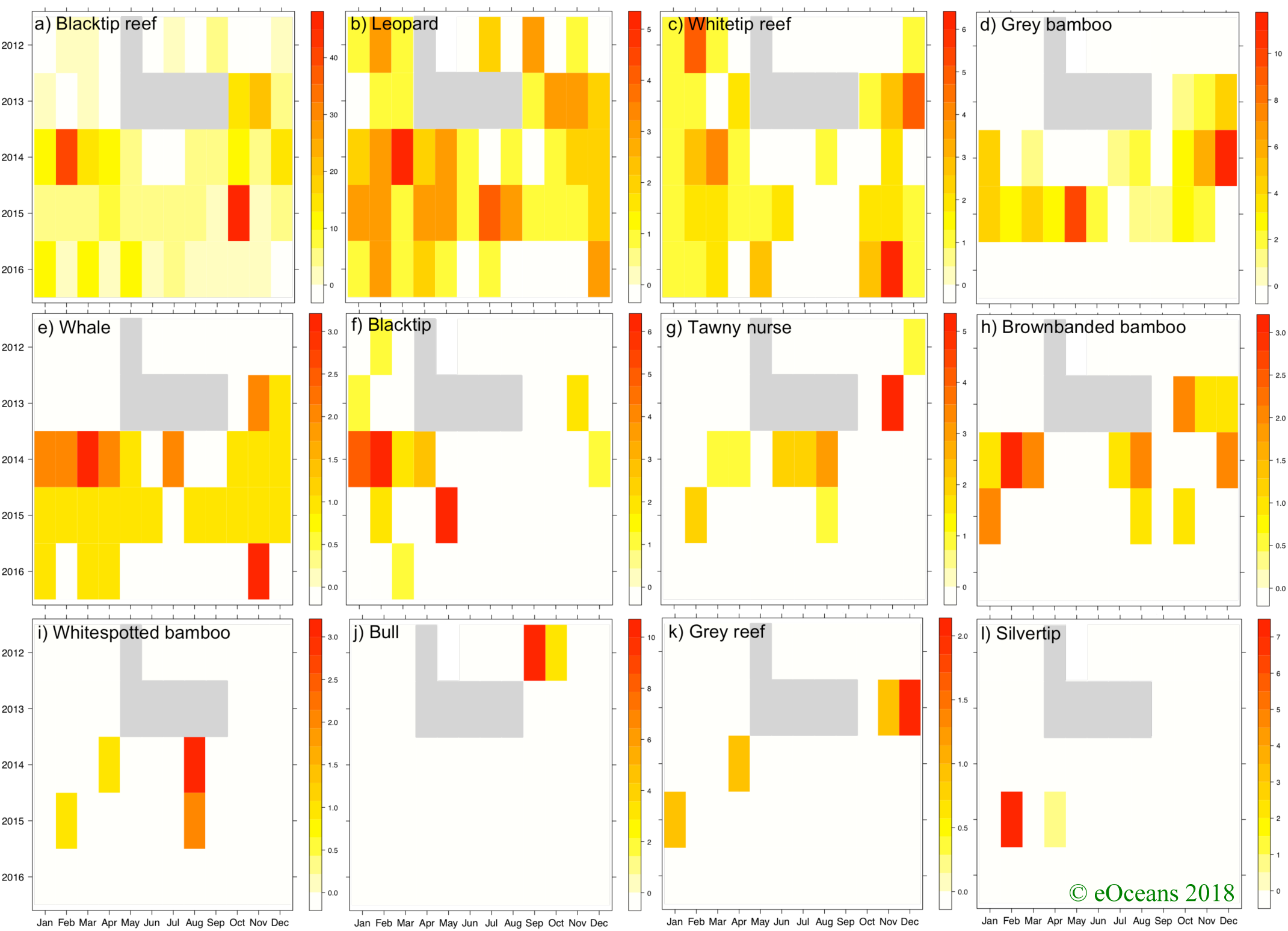
Temporal patterns of maximum school size by species. Each scale represents the largest school size observed for a species in that month. Grey boxes are months where sample effort was zero.

Sharks were observed on both coasts of Thailand throughout the study area, except in the northeastern sites near the Cambodian border (Figure 6). SF, maximum school size, and abundance varied by species (Table 1). Blacktip reef and leopard sharks were observed on the most number of sites, 47 and 43 sites representing 31% and 28% of all visited sites, respectively. On the sites where the species was observed (i.e., excluding zeros), mean SF was <10% (except the sites with silvertip sharks, which had a mean SF of 50% but were only encountered on two sites that were visited infrequently – 8 individuals in total, on two dives, where one site is named ‘Silvertip Bank’).

**Figure 6.**
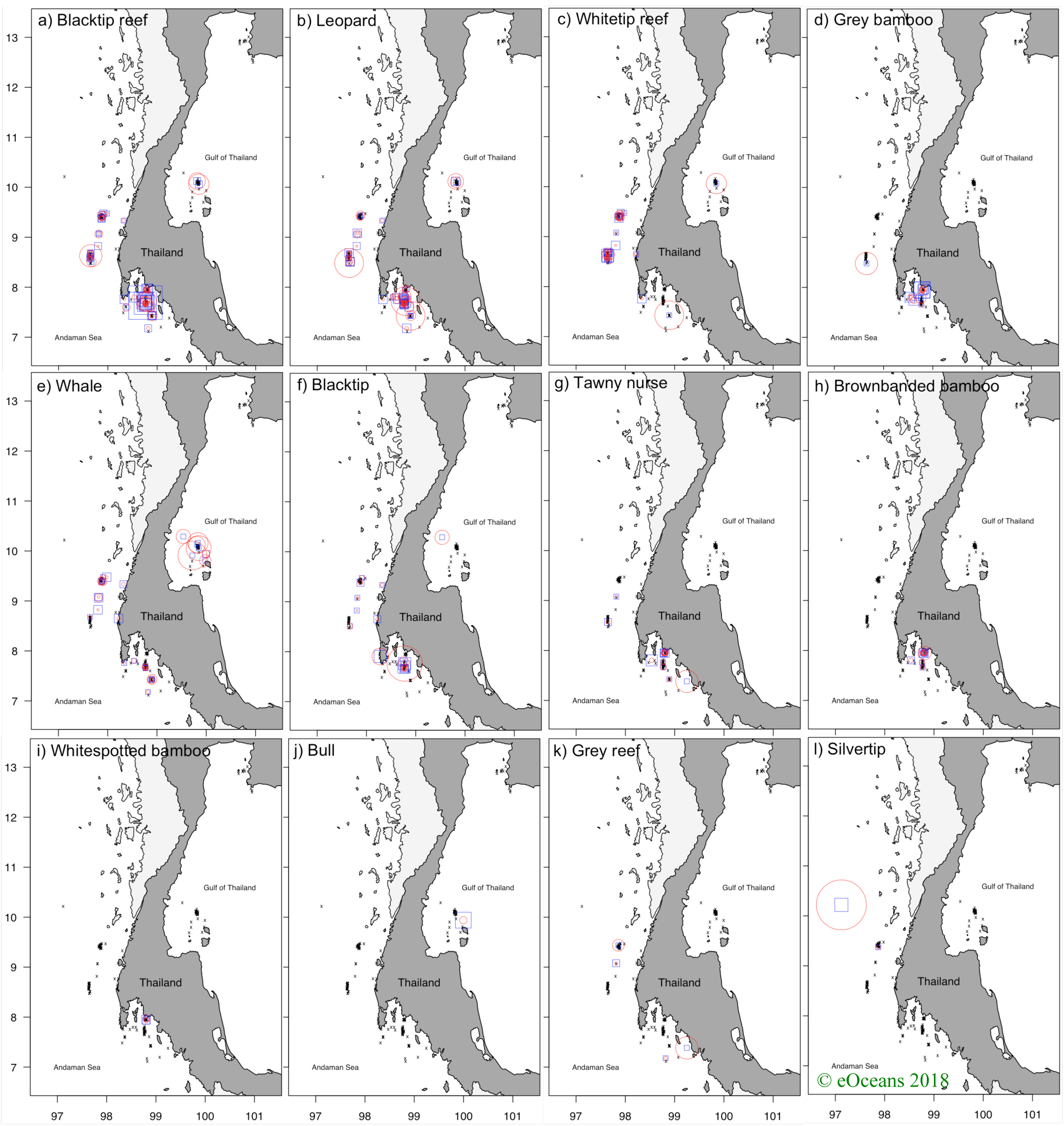
Spatial patterns of sighting frequency (red circles) and maximum number observed (blue square), where the size of each symbol represents the range of values (see Table 1 for range of values for each), and black x’s are sites with zero observations.

## I. DISCUSSION

Using an extensive database of underwater observations made by the dive community in Thailand, this study demonstrates the value of the eOceans event-based (i.e., where each dive is reported) monitoring platform for describing patterns of shark populations in coastal Thailand. This five year snapshot census of dive effort and sharks provides proxies of contemporary baselines for the ecosystem, and the social and economic value of these sites for the dive tourism industry, which may be used for monitoring future trends and informing policy. As well, the scale of the contributions, >9,000 dives on 153 sites over four years, shows the value of effective collaboration between scientists, non-governmental organizations, and the tourism industry.

### Observed patterns

Of the species and areas sampled, few have been described in detail before, especially in Thailand waters, which highlights the value and novelty of our findings. During this five-year study period, at least twelve shark species were encountered. According to the IUCN Red List Criteria (www.iucnredlist.org) two of these species are Vulnerable, eight are Near Threatened, and two are Endangered, including the whale shark (*Rhincodon typus*, Pierce and Norman, 2016) and the leopard shark (or zebra shark, *Stegostoma fasciatum*, Dudgeon et al., 2016) (Table 1). Spatial and temporal patterns have been described for some of these species elsewhere, such as seven studies documenting the movement patterns of blacktip reef sharks (Chapman et al., 2015, Table 1). However, there are few species for which spatiotemporal patterns of populations have been described (e.g., *Chiloscyllium griseum, Nebrius ferrugineus, Chiloscyllium punctatum, Chiloscyllium plagiosum*).

In some cases, the information gained from this study does support existing knowledge. For example, whale shark seasonal and annual trends in Thailand have been previously described by two different studies. Theberge and Dearden (2006) used divers’ observations collected by a single dive shop in the Andaman Sea, and Ward-Paige and Lotze (2011) surveyed dive instructors across the country to document spatial and temporal patterns of whale sharks (and other shark species). Both studies showed declines in sightings from the early 1990s to the early 2000s, with a total of only two to three individuals being encountered – exactly the maximum school size in the current study. Sighting frequency in Theberge and Dearden (2006) ranged from 8% (41/462) in 1992-1993 to <1% (2/339) in 2000-2001, which is also similar to the current study at 1.4% (133 sightings in 9,524 dives) sighting frequency in 2012-2016. Although the timelines do not overlap between these three studies, and the current study involved more observers and sites, the similarities suggest that various sampling strategies capture similar longer-term trends, and it optimistically suggests that previously documented declines may have been halted. Interestingly, however, is that whale sharks are considered seasonal, but were observed throughout the year, showing the value of continuous monitoring that includes zeros.

In other cases, our results differ from other studies. For example, one study from Thailand showed that divers commonly observe bull sharks on Chumphon Pinnacle, a site in the Gulf of Thailand (e.g., Brunnschweiler, 2009). In the early 2000s bull sharks were observed daily on this site, with up to 10 individuals being observed at a time (Ward-Paige and Lotze, 2011). They were so common that many interviewed dive instructors could consistently describe the daily and seasonal behaviour of these bull sharks, being deep in the morning and on top of the pinnacle in the afternoon (Ward-Paige and Lotze, 2011; CWP personal communication). However, in the current study, no bull sharks were observed on this site, and only one blacktip and two whale shark observations were made (Ward-Paige and Lotze, 2011; and verified with photo identification by CWP). This suggests that either effort was too low at this site to detect bull sharks (n = 12, various months), or that the population has changed (moved, or been lost) since these earlier studies, and further highlights the value and importance of sampling via eOceans into the future to infer trends.

Ecological studies from other regions may also provide insight into the metrics presented in the current study (i.e., occurrence, sighting frequency, school size). Blacktip reef sharks, for example, were the most commonly sighted species in the current study, and have been studied elsewhere. In other regions, blacktip reef sharks have been found to have long-term residency, and high site fidelity (Chin et al., 2013; Papastamatiou et al., 2010). In Thailand, where blacktip reef sharks were observed throughout the year, it is likely that many individuals, at least on a site-by-site basis, but perhaps also on nearby sites, are repeat observations of the same individuals. Additional support for this hypothesis comes from divers participating in this eOceans program who have been photographing and recording individual blacktip reef sharks, which show that some sharks are repeatedly observed at the same sites (e.g., orientalsea.com/ID-oversigt.htm). As well, blacktip reef sharks have been shown to undergo ontogenetic shifts in habitat selection, where adults used ledge habitats and pups used shallow waters, potentially as nursery areas (Papastamatiou et al., 2009). In the current study, four sites had high maximum school sizes (n = 11-45 individuals), all were near to, but not in, sheltered bays. These sites deserve further investigation to determine if they are, or are near to, essential habitats, such as pupping, mating, or nursery areas.

Sharks were observed on 59% of all sites, which is relatively high compared to other areas surveyed by divers that also had high nearby human populations (Ward-Paige et al., 2010b). However, some sites may have fewer sharks than could be expected. Six sites, for example, that are named after sharks had few shark sightings. Shark Fin Rock had no sharks, Whale Shark Wall had one blacktip reef shark, Shark Island had one whitetip reef and three blacktip reef sharks, Shark Point 2 had up to four grey bamboo and two leopard sharks, and Sharkfin Reef had up to three leopard, one whitetip reef and one blacktip reef shark. These sites could have been named for many reasons, other than the presence of sharks; however, they may indicate the loss of their namesakes. Ward-Paige and Lotze (2011), for example, found that many sites have lost whitetip reef sharks where they were once abundant. These discrepancies warrant further investigation.

### Implications

The patterns described herein may be viewed as a contemporary baseline against which future changes can be quantified. One of the primary goals of eOceans is to document and detect change in shark populations, and other animals, in response to anthropogenic impacts, environmental change, or as a result of protection or conservation measures. By collecting longitudinal observations of occurrence and maximum school size at a variety of sites, which includes participant effort and zeros (e.g., excursions where no sharks were observed), broad changes can be described. For example, using the sites described in the current study, including those where no sharks have been observed, we will monitor future changes in occurrence, maximum school size, and diversity through time. It is, however, possible for sharks to alter their distribution for many reasons, other than population change, and therefore any documented changes with eOceans data may be used to trigger additional studies.

By providing the scientific oversight to on-the-ground organizations, eOceans promotes collaboration and encourages broad-scale participation. eOceans brings the scientific expertise, including data collection, management, analysis and interpretation to a project like this, and the local leaders, like Shark Guardian, raise awareness about the project and encourage participation through their own missions. In this way, both objectives - data for science and opportunities for education and outreach - are met. These education and outreach opportunities should not be undervalued (Bonney et al., 2009) and soliciting observations of all divers across Thailand provided an unprecedented outreach avenue, creating one of the largest citizen science projects in Thailand to date. As such, Shark Guardian now delivers hundreds of presentations a year, thus reaching >20,000 people per year. During these presentations, eOceans data summaries and updates are provided, allowing participants to preview the data that has been collected thus far, to compare their own observations to that of the community. The eOceans program thus promotes personal connections between scientists, local organizations and businesses, and the general community of ocean divers and explorers. Thereby, using science as leverage for education and collaboration, and vice versa. Both organizations have also been discussing these results with the Thai government and other organizations working in the region (e.g., International Union for Conservation of Nature, IUCN) to ensure that these results are made available and are considered in policy decisions (e.g., Marine Protected Area Network design).

Increased participation in science by the broader community can also lead to higher acceptance rates of science and improved reception of recommended management and policy needs (Bonney et al., 2009; Cooper et al., 2007). As such, this community-led project in Thailand has both increased the collaborative network of recreational divers working together to census the marine environment, sharing the importance of data collection and gaining knowledge on shark and marine conservation, while collecting thousands of observations that can be used to fill important data gaps for scientific investigation.

As well, the alliance between the scientists at eOceans, the Shark Guardian team, and the network of dive shops, dive instructors, and divers presents a potentially powerful, scalable model that could be applied to other species or areas. Sharks are relatively conspicuous, with divers often repeatedly seeing the same species at a site, thus providing ample opportunity for photos to be taken and informed identifications to be made. However, many other species share similar qualities, such as rays, turtles, seahorses, jellyfish, and mammals, which eOceans also collect on in its online forms and will be used to describe spatial and temporal trends in further publications. As well, Thailand was an ideal location to test this type of research program since it has a large network of dive shops, readily available internet access, strong collaborative outreach leadership (in Shark Guardian), and frequently encountered sharks of various species. However, Thailand is not unique in many of these aspects, and it is expected that other areas or countries could similarly adopt this type of project, which could help fill important data gaps and provide ongoing monitoring.

eOceans data may also be useful for identifying priority conservation strategies, such as in the design of a National Plan of Action for Sharks (NPOA-Sharks; Fischer et al., 2012), “Hope Spots” (https://www.mission-blue.org/hope-spots/), or to design protected areas (e.g., to meet Aichi target 11, to protect 10% of ocean ecosystems in Marine Protected Areas) by identifying areas with the highest occurrence rates, diversity, or mating or nursery areas, and those that have high threats. Thailand has a long history of marine exploitation (Panjarat, 2008), is one of the top shark fishing nations (Fischer et al., 2012), has high human population, intensely modified coastlines, an abundance of boats, and few permanently protected areas, but sharks still remain in high enough abundance to be detected by divers. This cannot be said for many other regions of the world such as in the Caribbean, parts of Australia, and some outlying Pacific islands (Friedlander and DeMartini, 2002; Robbins et al., 2006; Sandin et al., 2008; Ward-Paige et al., 2010b). Nevertheless, previous studies suggest that sharks throughout the Indo-Pacific are vulnerable, have declined, and some, especially those dependent on coastal habitats, have essentially disappeared (Espinoza et al., 2014). Thus, although there is a paucity of data to delineate what has been lost, eOceans provides the data resource needed to begin to protect the sites where sharks remain in Thailand. Even if sites are only occasionally visited by rare or threatened species, for example, the sites may be important for their migration routes or essential life stages, and therefore deserve further investigation and consideration for protection.

### Caveats

There are a few caveats to consider with the results presented herein. This is an initial investigation of the data contributed by the recreational dive community in Thailand, and further analyses are warranted to provide more detailed insights on sharks and the participants in the region.

The majority of errors could be attributed to the design of the previous eOceans form and the use of common names, (e.g., *Stegostoma fasciatum* is Leopard shark locally, but Zebra shark in most identification books). However, there were 4 outliers (<0.2%) that were not corrected. Efforts are being made through education opportunities to ensure species are being correctly identified and reported, and validation at the time of submission into eOceans is recommended for future platform versions. As well, photos can now be submitted to eOceans, which may be used to evaluate identification errors. There remains the possibility that some species were misidentified, but given the analyses used, the effect is likely to be minimal. However, given that <0.2% of all submitted records contained outliers (later identified as mistaken entries), suggests high quality species identification and reporting amongst the participating dive community in Thailand.

Duplicate observations of the same shark could have been submitted, which would be an issue if the data were used incorrectly. However, with the analyses used here, all observations are considered duplicates of the same individuals at a site, which is why the maximum number observed is reported, and therefore underestimates the number of shark encounters. This assumption precludes some analyses, such as sightings per unit effort (e.g., as in Catch per unit effort). For many of these species that have relatively small home ranges, the maximum number observed may be a good representation of the total; however, movement studies may help to further define what would be reasonably considered a duplicate observation (e.g., species dependent observations made on the same day, month, year).

Divers also impact shark behaviour (Haskell et al., 2015; Vianna et al., 2014), which may skew observations (inflate or deflate bias; Ward-Paige et al., 2010a). This is an issue that needs to be considered with any sampling strategy. However, given the sheer numbers of divers in Thailand (it is one of the most heavily dived areas in the world), it is a unique place where fish are so used to divers that they come very close (CWP personal observation), suggesting that what is observed is at least a good representation of reality - similar to what could be expected by scientific divers’ observation. It is also one of the few places where sharks are regularly observed without using an attractant, again suggesting the impact of a divers’ presence may be relatively minimal. Regardless, the effects would be individual and species specific, and should not be dismissed, and therefore some rare or diver averse individuals may use sites when divers have left the site and may be underestimated.

Finally, despite relatively high sampling effort in comparison to many scientific studies, variability in effort is an issue to consider. Some sites were visited hundreds of times, thus increasing the chances of detecting sharks, and others only a handful of times, which may not be enough to detect sharks. As well, sites are selected based on preference of the divers, which may be related to skill level (e.g., shallow and protected) or interest (e.g., pinnacle), therefore skewing effort and precluding some analyses like habitat preference modeling. It is also possible that divers avoid sites with sharks, but this is unlikely given the economic potential of shark diving tourism (Cisneros-Montemayor et al., 2013; Vianna et al., 2012/1).

## I. CONCLUSION

Through effective collaboration with on-the-ground leadership teams in Thailand, eOceans gathered invaluable data to further knowledge on sharks, and fill important data gaps for these species in this region. Our results provided a unique perspective on sharks and the area covered by divers in Thailand, where little research has been conducted, thus providing the first glimpse into the seasonal, annual, and spatial distribution of a few species. Our results are particularly relevant because they provide data for the coastal zone of one of the top shark fishing nations, which has a long history of fishing. And despite this, it is remarkable that sharks remain in high enough abundance to be detected by divers, as the same cannot be said for other regions of the world. Our results could also help inform management and conservation strategies in the region. Since the sites investigated here are visited by the dive community, and are consequently likely some of the most economically valuable sites for ecotourism, there would be added value in investigating these sites to prioritize conservation actions.

## ACKNOWLEDGEMENTS

We are grateful to L Ward-Sing for essential on-the-ground and outreach support. We thank C Dudgeon, HK Lotze, and three anonymous reviewers for providing valuable comments on the manuscript. CWP thanks E Merritt and O MacKenzie for support. We appreciate all the Thailand dive shops who promoted the project, including: All 4 Diving, Amazing Phuket Adventures, Andaman Coral Divers, Aqua-vision, Big Blue Conservation, Big Blue Diving, Blue Guru, Blue View Divers, Hidden Depths Diving, IDC Thailand, IQ Dive, Khao Lak Explorer, Lanta Diver, Liquid Adventure, Master Divers, New Heaven Dive School, Projects Abroad, Redfish Diving, Rumble Fish Adventure, Scuba Cat, Scuba Fish, Sea Bees, Sea Dragon Dive Center, Sea Turtle Divers, Similan Diving Safaris, The Dive Inn, Wicked Diving. We are also thankful to each individual who carefully submitted their observations, including A Leeming, A Lemieux, A Roy, A Solomon, A Rugo, A Wolf, L Bindholt, B Norris, C Dudgeon, B Winkel, C Nicholls, C Lecky, Christa, Christoph, Clemmens, Cooper, Craher, L Cronin, D Sexton, D Weber, D Fernandez, D Tucker, Elaine, E Emili, E Loiseriou, F Lindgren, F Garz, H Macnee, A Higgins, I Landstrom, Janelle, Jasmin, J Wuzlon, J Zacho, J Bresseleers, J McGeachy, J Nolan, Jon, J Kettunen, K Bromfield, K Skellern, L Nelson, L Evans, M Sund, M Nelson, M Csere, N Bundy, Joe, P Ballard, Paurgell, P Hilgert, Phisher, P Macintosh, Rahul, Renatestein, Ric, Rob, Roupagoka, Rpaleg, Saffron, S Sherpa, S Luft, Settimo, S Hunert, Sinesara, S Arnold, S Srirachan, S Hinterberger, T Campbell Brown,T Sansal, Wymer, and many other contributors.

